# Global fMRI signal topography differs systematically across the lifespan

**DOI:** 10.1101/2022.07.27.501804

**Authors:** Jason S. Nomi, Danilo Bzdok, Jingwei Li, Taylor Bolt, Catie Chang, Salome Kornfeld, Zachary T. Goodman, B.T. Thomas Yeo, R. Nathan Spreng, Lucina Q. Uddin

## Abstract

The global signal (GS) in resting-state fMRI, known to contain artifacts and non-neuronal physiological signals, also contains important neural information related to individual state and trait characteristics. Here we show distinct linear and curvilinear lifespan patterns of GS topography in a cross-sectional lifespan sample, demonstrating its importance for consideration in studies of development and aging. Subcortical brain regions such as the thalamus and putamen show linear associations with the GS across the lifespan. The thalamus has stronger coupling in older-age individuals compared with younger-aged individuals, while the putamen has stronger coupling in younger individuals compared with older individuals. The subcortical nucleus basalis shows a u-shaped pattern similar to cortical regions within the lateral frontoparietal network and dorsal attention network, where coupling with the GS is stronger at early and old age, with weaker coupling in middle age. This differentiation in coupling strength between subcortical and cortical brain activity across the lifespan supports a dual-layer model of GS composition, where subcortical aspects of the GS are differentiated from cortical aspects of the GS. We find that these subcortical-cortical contributions to the GS depend strongly on the lifespan stage of individuals. Our findings demonstrate how neurobiological information within the GS differs across development and highlight the need to carefully consider whether or not to remove this signal when investigating age-related functional differences in the brain.

## Introduction

One of the biggest challenges in neuroscience is separating signal from noise (Uddin, 2020). In functional neuroimaging generally, and in human connectomics investigations using resting-state functional MRI (fMRI) data specifically, this challenge has been addressed with processing pipelines that mitigate artifacts known to obscure neural signals (Ciric et al., 2017; Parkes et al., 2018). The goal of these processing steps is to differentiate noise and relevant neural signals in fMRI data by removing physiological, hardware, and head motion-related signals to permit the discovery of underlying functional network architectures in the human brain. The “global signal” (GS) refers to the time series of signal intensity averaged across all voxels covering the brain, yielding one aggregate statistic per subject. The process of GS regression has been widely adopted as a robust method for attenuating noise due to cardiac and respiratory events and other confounding signals (Power et al, 2017). GS regression can also improve functional connectivity (FC) prediction of behavior (Li et al., 2019a). However, the GS is also an important component of brain function. Simultaneous fMRI-intracranial EEG studies in macaque monkeys demonstrate that gamma-band cortical electrical activity exhibits a positive correlation with BOLD changes across the entire cerebral cortex (Scholvinck, 2010) and unilateral suppression of the cholinergic basal forebrain causes changes in GS topography (Turchi et al., 2018). Simultaneous measurement of resting-state fMRI and calcium activity in awake rats has demonstrated significant correspondence between the GS measured non-invasively and neural spiking activity (Ma et al., 2020). Taken together, the emerging picture from these studies suggests that the GS contains relevant neural components, and does not simply represent noise in neuroscience investigations (Bolt et al., 2022; Li et al., 2019b).

The GS has also been shown to contain important information related to behavioral traits and intrinsic network organization in humans. We previously demonstrated that GS topography was related to a population axis of positive and negative life outcomes and psychological function, particularly weighted in frontoparietal executive control network regions, in a sample of over 1000 22-37 year old adults (Li et al., 2019b). Positive and negative life outcomes included measures of education, life satisfaction, cognitive flexibility, aggressive and internalizing behavior, alcohol abuse, and antisocial personality among others. More recently, we have shown that a dynamic spatiotemporal pattern that explains ∼20% of resting-state BOLD variance has a time series signature that is almost perfectly correlated (*r* = 0.97) with the GS (Bolt et al., 2022). This spatiotemporal pattern consists of negative cortex-wide BOLD amplitudes within somato-motor-visual (SMLV) complex, that then propagate toward cortical regions overlapping primarily with the frontoparietal network (FPN), but also with the default network (DN) and primary visual cortex (V1), followed by a spatiotemporal sequence with positive BOLD amplitudes with the same dynamics. These findings further suggest that the resting-state fMRI GS contains a rich source of important information relevant to large-scale brain network functional organization and individual differences in human cognition and behavior.

These results fit with the more recent conception of the GS as an important source of neural information, rather than being solely a source of noise. Accordingly, a recently developed dual-layer model of GS composition proposes that the GS represents two different layers of brain function (Zhang and Northoff, 2022). The first is a background subcortical-cortical layer where cortical activity is modulated by arousal and vigilance via subcortical regions such as the thalamus, basal forebrain, and midbrain. The second is a foreground cortico-cortical layer that is represented by network integration and segregation that is associated with cognitive states during rest and task. These two layers may operate in concert or independently to facilitate brain activity. This dual-layer model of the GS helps to reconcile the involvement of the GS in arousal, physiology, and cognition. However, it is currently unclear how subcortical and cortical brain activity contributing to the GS may differ across the lifespan.

Here we undertake a comprehensive assessment of age-related changes in spatial topography of brain regions associated with the GS across the lifespan. Despite the large amount of attention given to characterizing GS topography (for a review see: Ao et al., 2021) and the impact of GS regression on some of the most commonly deployed preprocessing pipelines (Ciric et al., 2017; Linkes et al., 2018; Power et al., 2017), the question of how age shapes the topography of the GS has not been carefully considered. Consequently, the extent that existing findings documenting lifespan changes in large-scale functional brain network configuration are potentially confounded with the differential implementation of GS regression across research groups is entirely unknown. We find distinct GS topography associations with age that were reliably present across multiple fMRI data preprocessing procedures. The findings suggest that the GS conveys neurobiologically meaningful information that changes over the course of human development, and developmental and aging studies that implement GS regression warrant careful reconsideration.

## Materials and Methods

### Subjects and fMRI data

A 10 minute resting-state fMRI scan was obtained from 601 subjects (6-85 years old; 240 males; **Supplemental Figure 1**) without a current Diagnostic and Statistical Manual of Mental Disorders (DSM) diagnosis from the Nathan Kline Institute (NKI) enhanced publicly available data repository (Nooner et al., 2012) (http://fcon_1000.projects.nitrc.org/indi/enhanced/). All participants provided written informed consent (written assent was obtained from minors and their legal guardian) for their data to be shared anonymously through the International Neuroimaging Data-Sharing Initiative (INDI) website (http://fcon_1000.projects.nitrc.org/).

Brain imaging was performed on a Siemens Trio 3.0T scanner that collected a T1 anatomical image and multiband (factor of 4) EPI sequenced resting-state fMRI data (2x2x2 mm, 40 interleaved slices, TR = 1.4s, TE = 30 ms, flip angle = 65°, FOV = 224 mm, 404 volumes). Participants were instructed to keep their eyes open and fixate on a cross centered on the screen. For quality control, we ensured that all participants had less than 0.5 mm average framewise displacement (FD). Linear regression revealed a significant linear FD-age association (β = 0.35, *p* = 1.23E-18) but no significant quadratic FD-age association (*p* = 0.9). Therefore, head motion was used as a nuisance covariate in all analyses.

### Preprocessing Pipelines

In order to account for non-neuronal artifacts and head motion, analyses were conducted across several preprocessing pipelines (MP - minimally preprocessed; CR - covariate regression; ICA-FIX; temporal ICA (tICA). Spatial ICA denoising, on which ICA-FIX is based, has been identified as one of the most effective tools for removing spatially structured noise artifacts from fMRI data (Ciric et al. 2018; Parkes et al., 2018). We also applied temporal ICA (tICA), which complements spatial ICA by removing temporally structured global noise (Glasser et al., 2018;). Collectively, these methods were investigated to ensure that non-neuronal artifacts were not driving the lifespan GS topography associations. These preprocessing pipelines demonstrate that GS topography associations with age are robust to a range of widely-used denoising procedures.

### Minimally Preprocessed (MP) Pipeline

All resting-state fMRI data were preprocessed using FSL, AFNI, and SPM functions through DPARSF-A in DPABI (Yan & Zang 2010). The first five images were removed to allow the MRI signal to reach equilibrium. Next, resting-state fMRI data were despiked using AFNI 3dDespike, realigned and normalized with DPARSF-A into 3mm MNI space using a priori SPM EPI templates, smoothed using AFNI 3dBlurToFWHM (6mm), and bandpass filtered using DPARSF-A (0.01 - 0.1 Hz).

### Covariate Regression (CR) Pipeline

After smoothing, DPARSF-A was used to calculate and regress out nuisance variables for covariate regression (CR) consisting of the Friston 24 motion parameters (six rigid-body head motion parameters, the previous time point for all six parameters, and the 12 squared derivatives; Friston et al., 1996), white matter time-series and cerebral spinal fluid time-series (using DPABI default masks), and a linear detrend. Finally, the data were bandpass filtered.

### ICA-FIX Denoising

Subject-level spatial ICA denoising (Griffanti et al., 2014) was conducted using ICA-FIX on minimally preprocessed data that was smoothed, but not subjected to covariate regression or bandpass filtering. The ICA-FIX classifier was trained on hand-classified independent components separated into noise and non-noise categories on data from 24 subjects (randomly sampled by choosing subjects separated by ∼10 years of age, and also choosing subjects with small and large amounts of head motion). Noise and non-noise components were classified by visual inspection using component maps, time-series, and power spectra (Griffanti et al., 2017). The resulting component classifications were then fed into FMIRB’s ICA-FIX classification algorithm (Salimi-Khorshidi et al., 2014) to automatically classify noise and non-noise components from individual subject data. Next, components classified as noise were regressed out of the data. Finally, the Friston 24 motion parameters and a linear trend were regressed out of the data, before a bandpass filter was applied.

### Temporal Independent Component Analysis (tICA) Denoising

The temporal ICA (tICA) pipeline was conducted using the FastICA algorithm in Python (https://scikit-learn.org/stable/modules/generated/sklearn.decomposition.FastICA.html). The tICA first conducted a group spatial ICA (sICA) on all 601 resting-state scans producing 125 independent components ( Glasser et al., 2018; Smith et al., 2012; **Supplemental Figures 2-6**). Classification of noise and non-noise components were conducted according to the procedure detailed in the ICA-FIX pipeline. Thirty-nine sICA components classified as noise were then regressed out of the remaining 86 non-noise sICA component time-courses. The cleaned time courses from the sICA were then concatenated across subjects to produce 86 time-courses, each with 239,799 TRs (399 TRs x 601 subjects). These concatenated cleaned sICA time-courses and representative group component spatial maps were then subjected to a tICA that produced 75 tICA time-courses and the associated group tICA spatial component maps (**Supplemental Figures 7 and 8**). Temporal ICA components can be classified just as spatial ICA components with visual identification of network activity and noise activity (Glasser et al., 2018). Nineteen of the 75 tICA components were identified as noise **(Supplemental Figure 9**). These components consisted of anti-correlated activity within the brain stem (e.g., TC 2, TC 17, TC 18) and striped banding representing head motion (e.g., TC 14, TC 28). Temporal ICA component 72 (**Supplemental Figure 9**) showed a general overall negativity across the cortex with little anti-correlation. Such tICA components in previous work have been proposed to represent a global component thought to be associated with the noise aspects of the GS (Glasser et al., 2018; Smith et al., 2012).

Finally, the time-series from the 19 noise group tICA components, the 39 group sICA noise components, the Friston 24 motion parameters, and a linear detrend were regressed out of each subject’s resting-state data before a bandpass filter was applied. More conservative and liberal classifications of tICA noise components (that did and did not include tICA component 72 (**Supplemental Figure 9**) produced similar results to the original 19 noise-component classification presented here. This shows that the noise and non-noise classification criteria of tICA components did not influence the pattern of results presented here.

### Scrubbing

Frames where FD exceeded 0.5 mm (Power et al., 2012) were not included in the regression model (average number of scrubbed frames per subject was 31.43 of 399 TRs (12.70%).

### Global Signal Topography and the General Linear Model

The GS was calculated as the mean time-series of all gray matter voxels within a SPM gray matter probability mask thresholded at 20%. Previous research has shown that the GS calculated across all voxels (white matter, CSF, etc.) in the brain compared to the GS calculated across only gray matter voxels in the brain are nearly identical (Glasser et al., 2018; Li et al., 2019) making it unlikely that the current methodology influenced the results. Linear regression between the GS time-series and the time-series of each voxel produced whole-brain voxel-wise beta maps. Frames where FD (Power et al., 2012) exceeded 0.5 mm were not included in the regression model. Next, each individual subject’s beta map was converted to z-statistics. Two general linear models (GLM) were then run in FSL using the whole-brain voxel-wise beta maps for all participants as the dependent variable (DV). The first GLM included linear age, mean FD, and sex as independent variables (IV) while the second GLM included linear age, quadratic age, mean FD, and sex as IVs. Age was the IV of interest within the first model and quadratic age was the IV of interest within the second model. The resulting group spatial maps were thresholded in FSL (voxel-wise uncorrected at *p* < 0.001 and cluster-wise corrected at *p* < 0.05) using Gaussian Random Field (GRF) theory. The two GLMs were run across all four preprocessing pipelines.

## Results

### Global Signal Topography across the Lifespan

Global signal topography maps showed increased coupling between the GS and visual, frontal, and sensorimotor brain regions (**Figure 1**). GLM results show that GS topography has distinct cross-sectional associations over the lifespan across subcortical and cortical brain regions. For subcortical brain regions, the thalamus shows a strong positive linear relationship with age, where coupling between the thalamus and GS time-series (scatterplot represents voxels within the thalamus thresholded at z > 5 for presentation purposes) increased across the lifespan. (**Figure 2**). The nucleus basalis (-18, -2, -12) showed a positive quadratic effect where coupling with the GS was weakest during middle age but stronger in young and old age. Finally, the putamen showed a negative linear relationship with age where coupling with the GS is strongest in early age but weakest during old age. These results demonstrate that subcortical regions involved in arousal and vigilance have distinct age-dependent cross-sectional associations with the GS across the lifespan.

**Figure 1:**
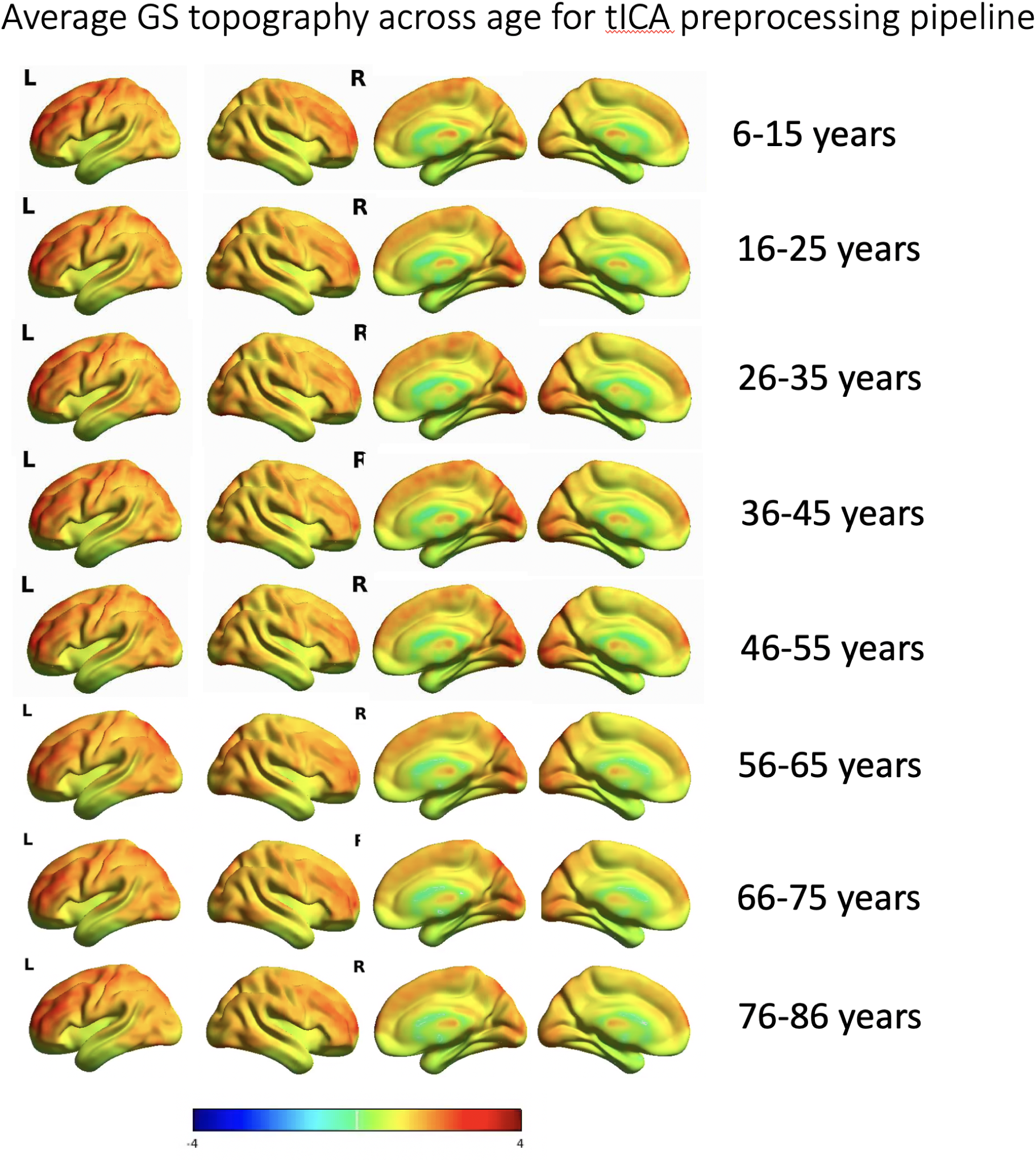
Average global signal topography across ten year age groups. Increased coupling between the GS with visual, sensorimotor, and prefrontal cortical regions are found across each age group.

**Figure 2:**
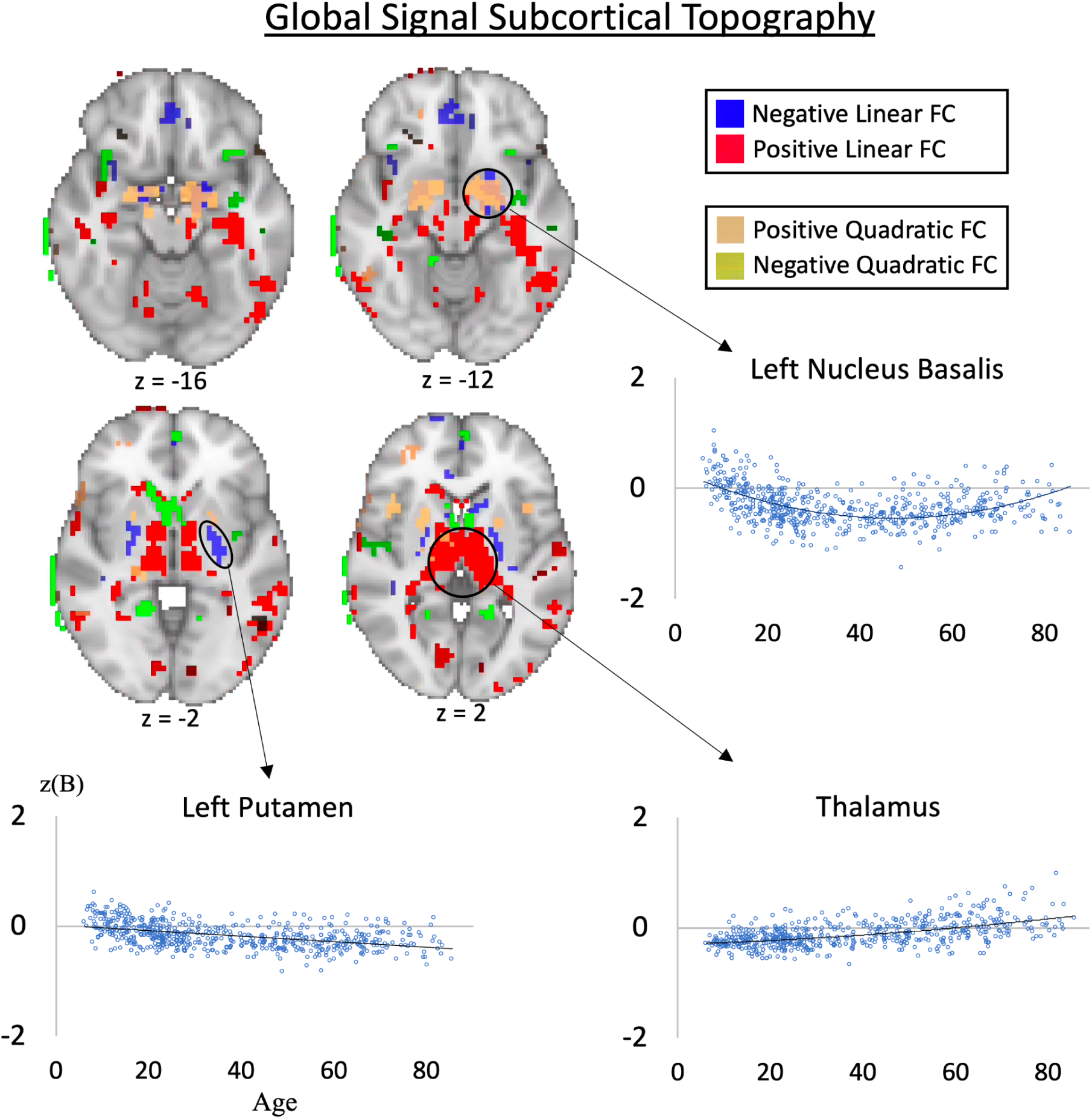
Group spatial maps showing subcortical associations between the global signal and voxel-wise functional connectivity strength across the lifespan (*p* < 0.001 voxel-wise uncorrected and *p* < 0.05 cluster-wise corrected). Negative quadratic associations show that the global signal has weaker associations with the nucleus basalis in middle aged individuals compared with younger and older individuals. The thalamus shows a positive linear association, where coupling with the global signal is stronger in older individuals compared with younger individuals. The putamen shows a negative linear association where coupling with the global signal is stronger in younger individuals compared with older individuals. The z-scored unstandardized beta is on the y-axis and age is on the x-axis.

For cortical brain regions, the lateral frontoparietal control network (parietal cortex overlapping with Schaefer ROI 333) (17 Network 400 ROI parcellation; Schaefer et al., 2018), dorsal attention network (inferior temporal cortex overlapping with Schaefer ROI 271, 272; FEF overlapping with Schaefer ROI 61, 261) showed a quadratic association with the GS where network nodes coupled with the GS are strongest at early (< 20 years) and later (> 60 years) periods of life, and the weakest in middle age (**Figure 3**). These results demonstrate distinct age-dependent large-scale network associations with the GS in networks related to external attention and cognition such as the control network and dorsal attention network.

**Figure 3:**
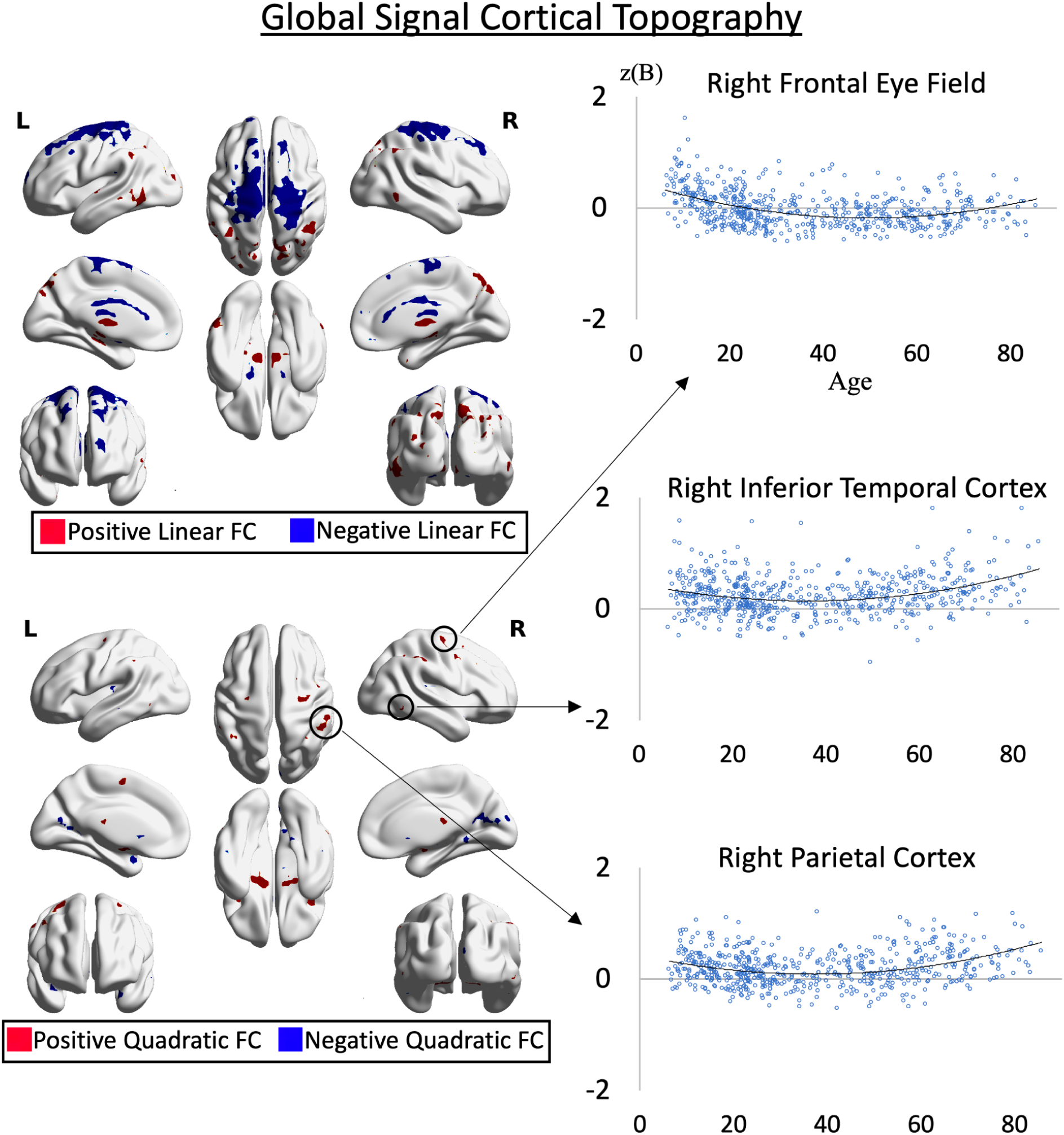
Group spatial maps showing cortical associations between the global signal and voxel-wise functional connectivity strength across the lifespan (*p* < 0.001 voxel-wise uncorrected and *p* < 0.05 cluster-wise corrected). Positive quadratic associations show that the global signal has stronger associations with regions of the lateral frontoparietal (parietal cortex) and dorsal attention networks (frontal eye fields and inferior temporal cortex) in younger and older individuals compared with middle aged individuals. The z-scored unstandardized beta is on the y-axis and age is on the x-axis.

### Preprocessing Pipelines and Imaging Artifacts

Both linear and quadratic lifespan coupling between GS topography and age were largely unaffected by preprocessing choices and produced the same linear and quadratic cross-sectional age effects as the main analysis (**Supplemental Figures 10 and 11**). In order to ensure that different preprocessing pipelines did change the composition of the GS time-series while leaving GS topography cross-sectional effects with age generally unaffected, within-subject temporal correlations between GS time-series across different preprocessing pipelines were calculated.

The within-subject average GS time-series showed a strong temporal correlation across all preprocessing pipelines (*r*s = 0.73 - 0.96) (**Figure 4**). The temporal correlation of the GS time-series between the FIX pipeline and the tICA pipeline was *r* = 0.76. The lower correlation between the FIX and tICA pipelines shows that the addition of tICA denoising has a large influence on the composition of the global signal, demonstrating that a large amount of variance was removed (*r* = 0.76; R^2^ = 58% variance explained). This demonstrated the effectiveness of tICA denoising in removing spatially and temporal structured fMRI noise within resting-state data that contribute to the GS, and also demonstrates the robustness of the current GS topography-age effects.

**Figure 4:**
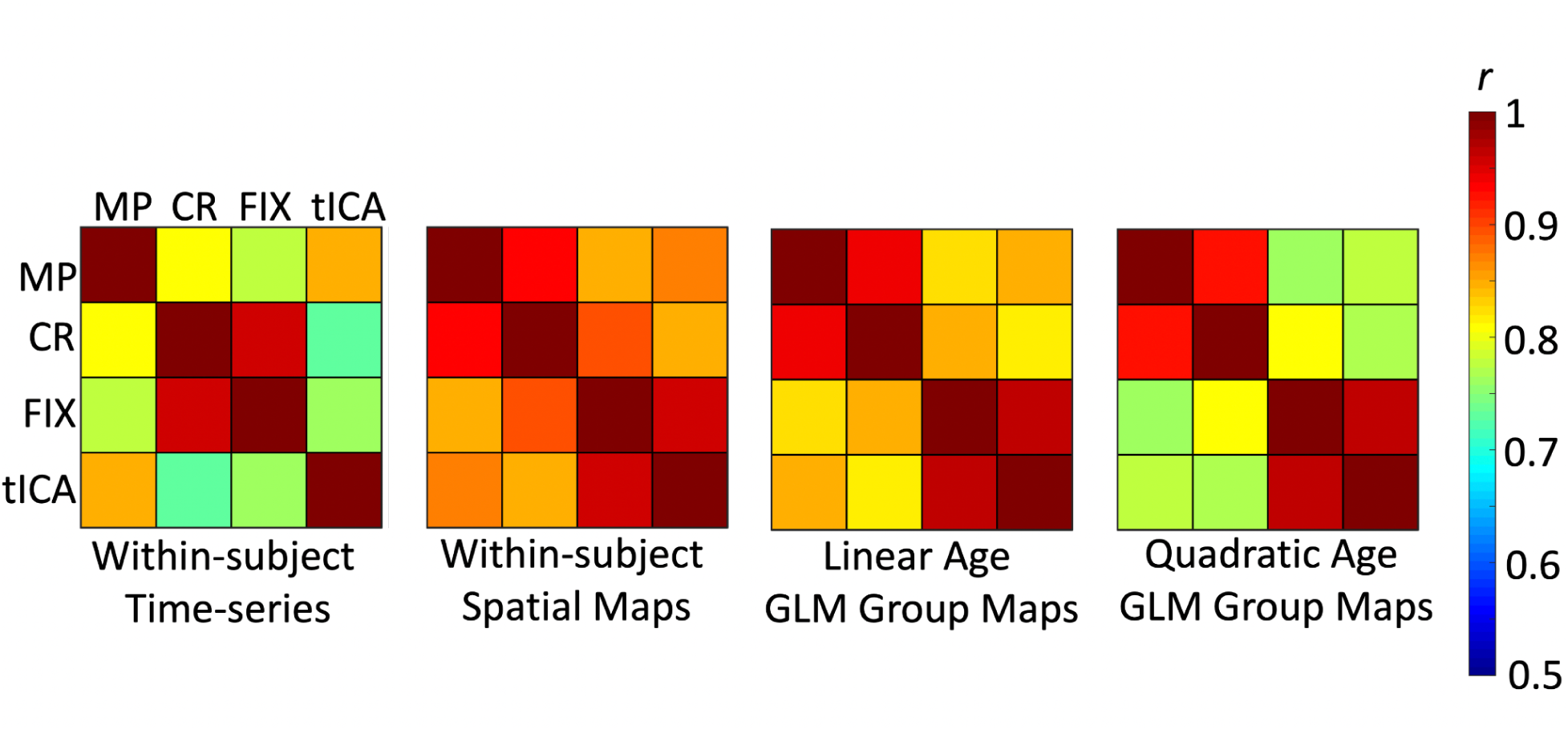
Temporal and spatial correlations across preprocessing pipelines. Within-subject time-series temporal correlations and within-subject maps spatial correlations represent the influence of preprocessing pipelines across subjects without considering the influence of age. Linear age GLM and quadratic age GLM effects represent the influence of preprocessing pipelines across group GLM effects for linear age and quadratic age. All matrices show strong associations across preprocessing pipelines (*r’*s > 0.73). MP = minimally preprocessed; CR = covariate regression; FIX = ICA-FIX, tICA = temporal ICA.

Despite the large amount of variance removed from tICA denoising compared with ICA-FIX in the GS time-series, the within-subject GS topography spatial map correlations showed little changes between the FIX pipeline and the tICA denoising pipeline (*r* = 0.96). This demonstrates that spatially and temporally structured noise do not significantly contribute to GS topography composition (**Figure 4**); if spatially and temporally structured noise did have an influence on GS topography, the within-subject spatial map correlation between the FIX and tICA pipelines should be much lower. Thus, although tICA denoising results in a quantifiable change in the GS time-series composition, GS topography remains unaffected.

Group level spatial correlations between the voxel-wise GLM linear and quadratic spatial map outputs were used to quantify the influence of preprocessing pipeline on lifespan age effects. These results showed that spatial patterns of linear (*r*s = 0.81 - 0.97) and quadratic (*r*s = 0.77 - 0.97) GS topography GLM results for age effects are similar across preprocessing pipelines (**Figure 4**). Thus, while the tICA preprocessing pipeline removes a significant amount of GS time-series variability, the within-subject GS spatial topography and group GLM GS spatial topography cross-sectional age effects remain relatively stable.

### Head Motion Considerations

To further ensure that head motion was not driving GS topography changes across the lifespan, we used a multivariate partial least squares analysis (PLS) implemented in Matlab (McIntosh, Chau, & Protzner, 2004) to identify the spatial relationship between head motion and GS coupling strength across the GS topography spatial maps from the tICA preprocessing pipeline. PLS maximizes the covariance among voxels with a behavioral variable of interest that is represented by a latent variable (LV) (**Supplemental Figure 12**). This LV represents the multivariate whole-brain voxel-wise spatial relationship between head motion (as measured by FD) and GS topography across subjects, and is differentiated from random noise using permutation testing (5000 permutations, *p* < 0.05). Each voxel within the LV is subjected to bootstrap estimation of standard errors (5000 bootstraps; approximates a p value of < 0.001) to determine if the voxel score is reliably different from zero. Each subject is then assigned a “brain score” that represents how strongly that subject’s data is represented within the LV. Each subject’s brain score was then used as an additional nuisance regressor in the linear and quadratic GLMs assessing the association between GS topography magnitude and age, in addition to the average FD head motion and sex regressors from the original analyses. The results were unchanged from the main analyses (**Supplemental Figure 13**), demonstrating that the influence of head motion on GS topography is spatially distinct from the influence of age on GS topography.

## Discussion

Global signal regression is a widely used fMRI preprocessing step, yet this practice remains one of the most controversial topics in network neuroscience (Liu et al., 2017; Murphy & Fox, 2007; Uddin, 2020). By providing a whole-brain metric of average brain activation (i.e., the GS) coupled with individual voxel activation, GS topography represents a unique representation of intrinsic brain organization related to trait behavior, task states, and clinical diagnosis (Ao et al., 2021; Li et al., 2019b). Our results show that coupling between brain regions and the GS depends on lifespan stage and spatial location in the brain. We also find that GS topography associations across the lifespan are stable across multiple preprocessing pipelines across the lifespan, demonstrating support for GS topography as a useful way of characterizing overall brain activity and connectivity related to development. Our results demonstrate the utility of GS topography in characterizing brain organization across the human lifespan and also suggest that careful consideration of GS regression is warranted when age-related FC effects are of interest.

The current study shows quadratic patterns of coupling between the GS and network nodes within the lateral frontoparietal control network and dorsal attention network. Relative to middle age, these two networks show stronger associations with the GS at early (< 20 years) and later periods (> 60 years) of life. The opposite pattern emerges in the medial prefrontal cortex, caudate, and lower-level visual cortices, where the association with the GS is weakest at early and later periods of life, with the strongest association with the GS in middle age. These quadratic associations are similar to lifespan trajectories of functional connections that typically show a curvilinear pattern of network development, where within-network coupling increases while between-network coupling decreases until middle age (Fair et al., 2009). After middle age, within-network coupling increases while within-network decreases (Betzel et al., 2014; Chan et al., 2014; Vij et al., 2018). The trajectory of the lateral frontoparietal network also closely resembles executive function performance across the lifespan, where performance peaks in the 3rd and 4th decade of life before dropping off in old age (Ferguson, Brunsdon, & Bradford, 2021). Taken together, curvilinear trajectories of network integration and segregation, as well as executive function behavioral performance show similar curvilinear trajectories as the lateral frontoparietal control network. Thus, our results support a dual-layer model of GS composition that demonstrates linear cross-sectional changes within the thalamus and sensorimotor regions as part of the background arousal layer against quadratic cross-sectional lateral frontoparietal control network changes in the foreground cortico-cortical cognition layer.

The current study showed that subcortical regions have distinct coupling patterns with the GS across the lifespan. The thalamus presented with stronger coupling with the GS across the lifespan. The thalamus has been identified as an integral initiating and mediating force of arousal and vigilance in the brain, as well as facilitating shifts in connectivity, activity, and network topology (Shine et al., 2023). Stronger thalamic coupling with the GS across age could indicate that portion of the GS related to arousal becomes increasingly important across the lifespan. On the other hand, the thalamus has been shown to play an important role in aging and cognition in task-fMRI studies (Goldstone, Mayhew, Hale et al., 2018). Thus, it is possible that the thalamus plays an integrative role in both vigilance and cognitive processes across the lifespan in the context of its role in GS composition.

The nucleus basalis presented with stronger coupling with the GS at early and late periods of life compared to middle age. Previous research has demonstrated that deactivation of the nucleus basalis via chemical intervention in macaques modulates the BOLD GS in the ipsilateral hemisphere (Turchi et al. 2018). This suggests a causal role of the nucleus basalis in cortical BOLD activity. The current study shows that the nucleus basalis has the same u-shaped pattern of coupling with the lateral frontoparietal and dorsal attention networks. Within the context of the current study, this may suggest that the contribution of nucleus basalis activity to the GS coordinates activity within the lateral frontoparietal control and dorsal attention networks. However, it is not possible to determine the causal direction of this relationship across the lifespan as in vivo manipulation of the nucleus basalis is not possible to conduct safely in humans.

The results are in accord with the dual-layer model of GS composition (Zhang & Northoff, 2022) that proposes a subcortical-cortical background layer associated with arousal and vigilance via the thalamus and basal forebrain (Liu et al., 2018) and a cortico-cortical foreground layer associated with network organization and cognitive rest-task states (Zhang et al., 2021).

Within the context of the current study we find that subcortical and cortical contributions to the GS vary across the lifespan. In early life, the GS shows stronger coupling with the putamen, caudate nucleus, lateral frontoparietal control network, and the dorsal attention network. In middle age, the coupling between these regions and the GS is weakest with only the thalamus showing increased coupling with the GS. Finally, in old age, all subcortical and cortical regions show strong coupling with the GS besides the putamen which shows its weakest coupling with the GS in older age. These differing patterns of coupling across the lifespan suggest that the contribution of various brain regions to the GS change across the lifespan. The changing composition of brain activity contributing to the GS across the lifespan may be an attempt at optimizing arousal and vigilance processes of the background subcortical-cortical layer with cognitive processes of the cortical foreground layer. Future studies will need to examine the mechanisms driving these associations such as identifying how differing levels of brain activity may be driving relationships with the GS time-series and in turn, influencing GS topography.

Although previous research suggests that temporal ICA denoising effectively removes structured global artifacts such as head motion, respiration, and cardiac events from resting-state fMRI data (Glasser et al., 2018), it is still possible that unstructured spatial and temporal noise has an influence on GS topography and its lifespan associations. Importantly however, the significant change of GS composition between FIX and tICA preprocessing pipelines, combined with the fact that the spatial location of age-FC GS topography effects remained virtually unchanged, suggests that GS topography is somewhat robust to such artifacts. Additionally, it is unclear if one would want to completely remove respiration and cardiac associated with neural function as they play an important role in the dual-layer model of GS composition (Zhang & Northoff, 2022) as they are intricately linked with the global signal (Bolt et al., 2023). These factors along with previous research showing that the GS is strongly associated with brain network activity (Gotts et al., 2020) and behavioral traits (Li et al., 2019) show how GS topography can be of further interest to neuroscientists as a biologically important aspect of brain function.

The systematic trajectories of GS topography across the lifespan and difference in age-FC strength relationships when using GS regression makes interpretation of studies using GS regression and age as a variable of interest more complex. As the age range of the sample increases, there is a greater possibility that different brain regions and networks will be influenced by GS regression. For example, GS regression may have a greater influence on the lateral frontoparietal network in younger and older age samples compared with middle age samples. Additionally, GS regression on young individuals may not influence thalamic and occipital cortex activity as much as GS regression on older adults. Thus, GS regression may have system-specific implications in categorical and dimensional fMRI age investigations. Further compounding these issues is that it is unknown if GS regression will be beneficial or detrimental for identifying cognition related brain activity. That is, it is not possible to determine if the GS is driving activity in specific networks, or if specific network activity is driving the global signal in an age-dependent manner. GS regression would be beneficial in the former case, but detrimental in the latter. Currently, the underlying physiological and neuronal contributions to the global signal remain unknown.

In conclusion, we show that age is significantly associated with the spatial topography of the GS in resting-state fMRI data. The thalamus and sensorimotor regions show distinct linear cross-sectional lifespan patterns compared with the quadratic lifespan patterns found for the lateral frontoparietal control network. Our results support a dual-layer view of the GS where composition of the GS may include a subcortical-cortical background layer modulating arousal via the thalamus and a cortico-cortical foreground layer modulating cognition via the lateral frontoparietal network that diverge as linear and quadratic effects across the lifespan. Due to the importance and unabated controversy over GS regression, researchers should be cautious when considering the implications of its application. As the field of fMRI keeps maturing, understanding how GS regression may help or hinder statistical analyses, and potentially mask true age-related FC effects, will continue to be of paramount importance.

## Data and Code Availability

All data are available for download from the Enhanced Nathan Kline Institute - Rockland Sample data repository (http://fcon_1000.projects.nitrc.org/indi/enhanced/).

Code used in analyses available on Github: https://github.com/jasonSnomi/LifespanGlobalSignalTopography

## Author Contributions

J.S.N. conceived of the project, helped with data preprocessing, conducted all analyses, and wrote the manuscript. D.B. and L.Q.U. conceived of the project, consulted on analyses, and helped write the manuscript. J.L., B.T.T.Y, and C.C. consulted on the analyses and helped write the manuscript. S.K. and Z.T.G. helped to download, sort, and preprocess the resting-state data. T.B. and R.N.S. contributed to the data analysis and helped to write the manuscript.

## Acknowledgements

BTTY is supported by the NUS Yong Loo Lin School of Medicine (NUHSRO/2020/124/TMR/LOA), the Singapore National Medical Research Council (NMRC) LCG (OFLCG19May-0035), NMRC CTG-IIT (CTGIIT23jan-0001), NMRC STaR (STaR20nov-0003), Singapore Ministry of Health (MOH) Centre Grant (CG21APR1009), the Temasek Foundation (TF2223-IMH-01), and the United States National Institutes of Health (R01MH120080 & R01MH133334). Any opinions, findings and conclusions or recommendations expressed in this material are those of the authors and do not reflect the views of the Singapore NRF, NMRC, MOH or Temasek Foundation.

---

**Supplemental Figure 1:**
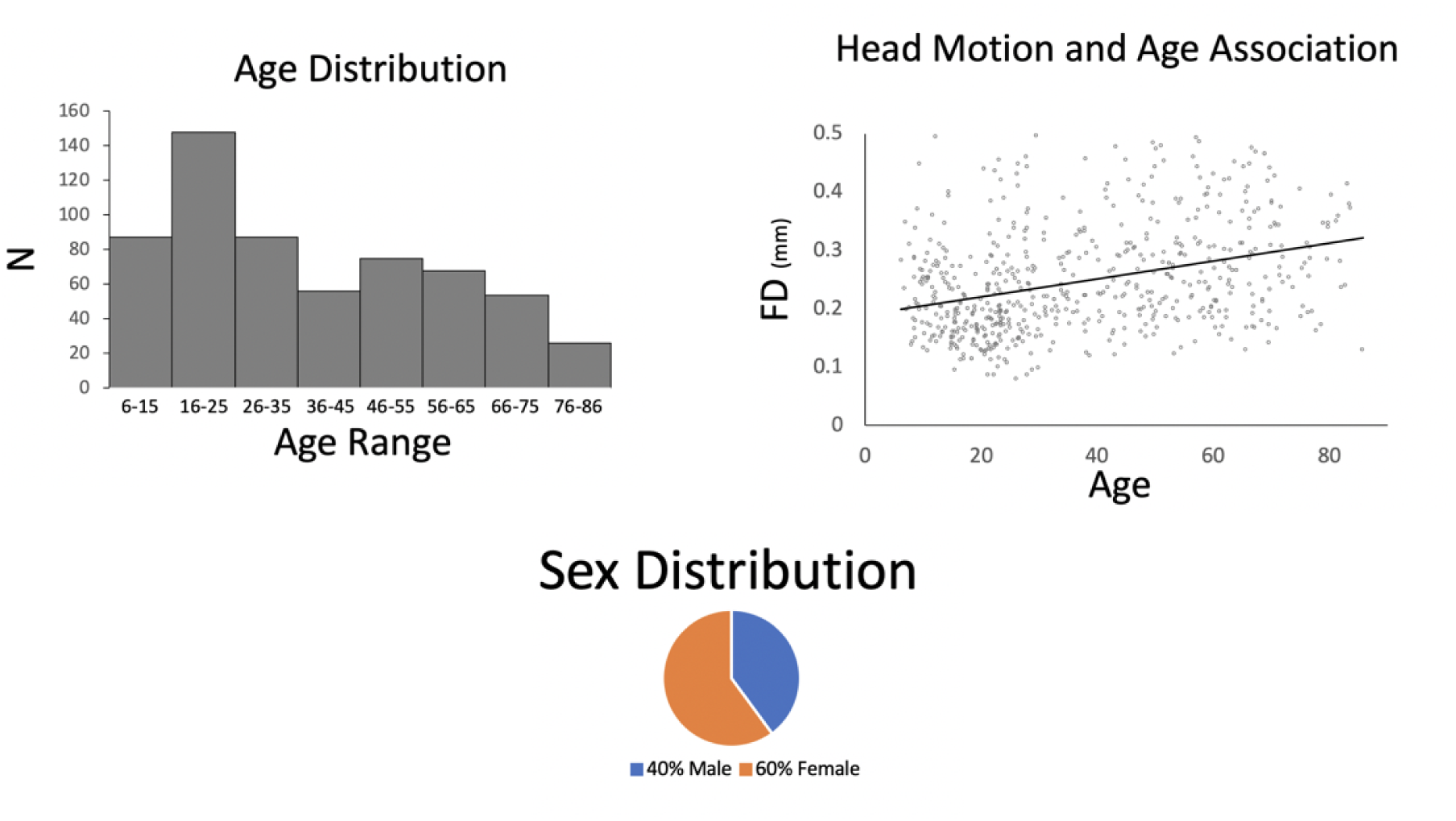
Age and sex distribution and the relationship between head motion and age.

**Supplemental Figure 2:**
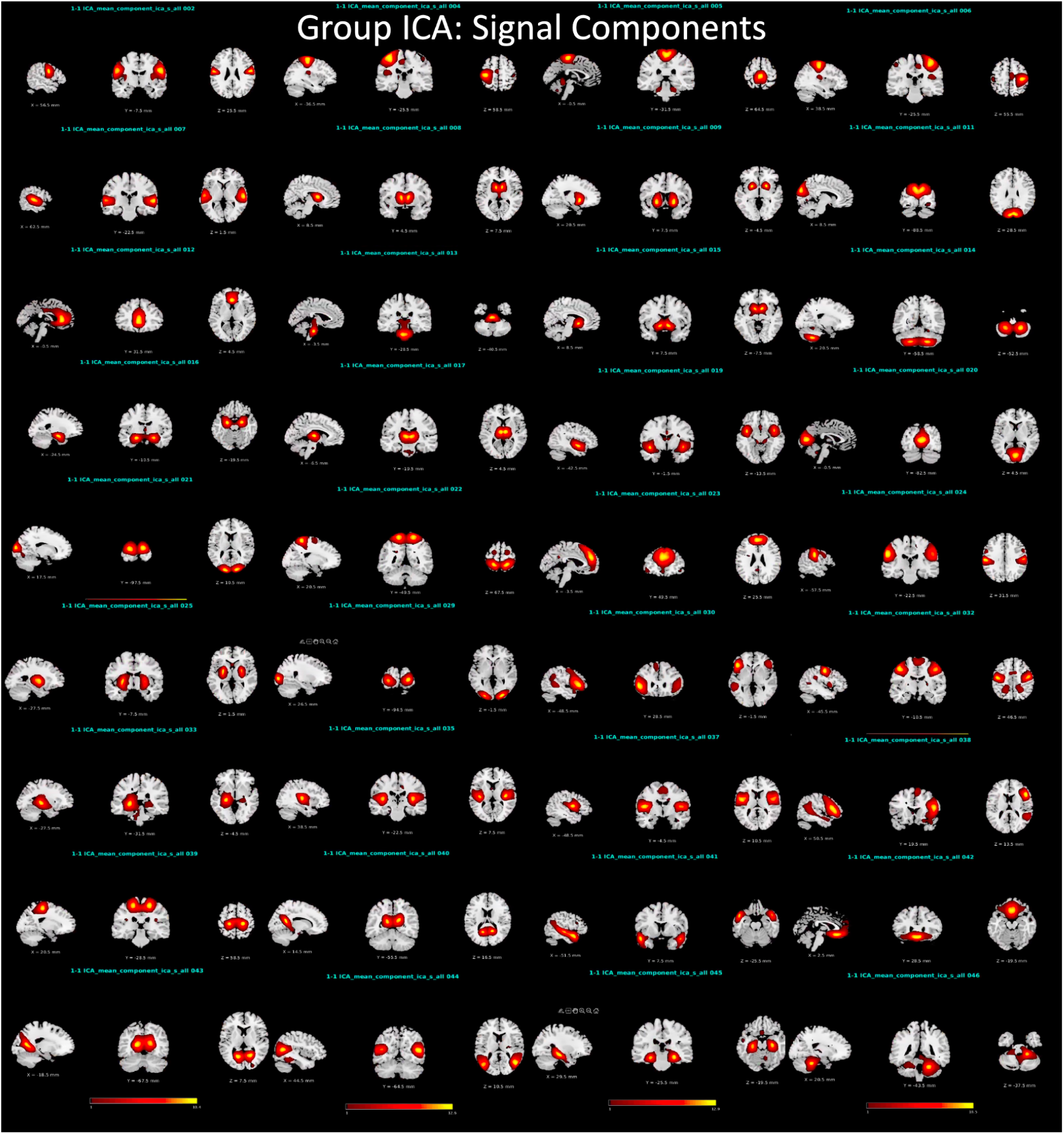
Group spatial ICA components classified as signal.

**Supplemental Figure 3:**
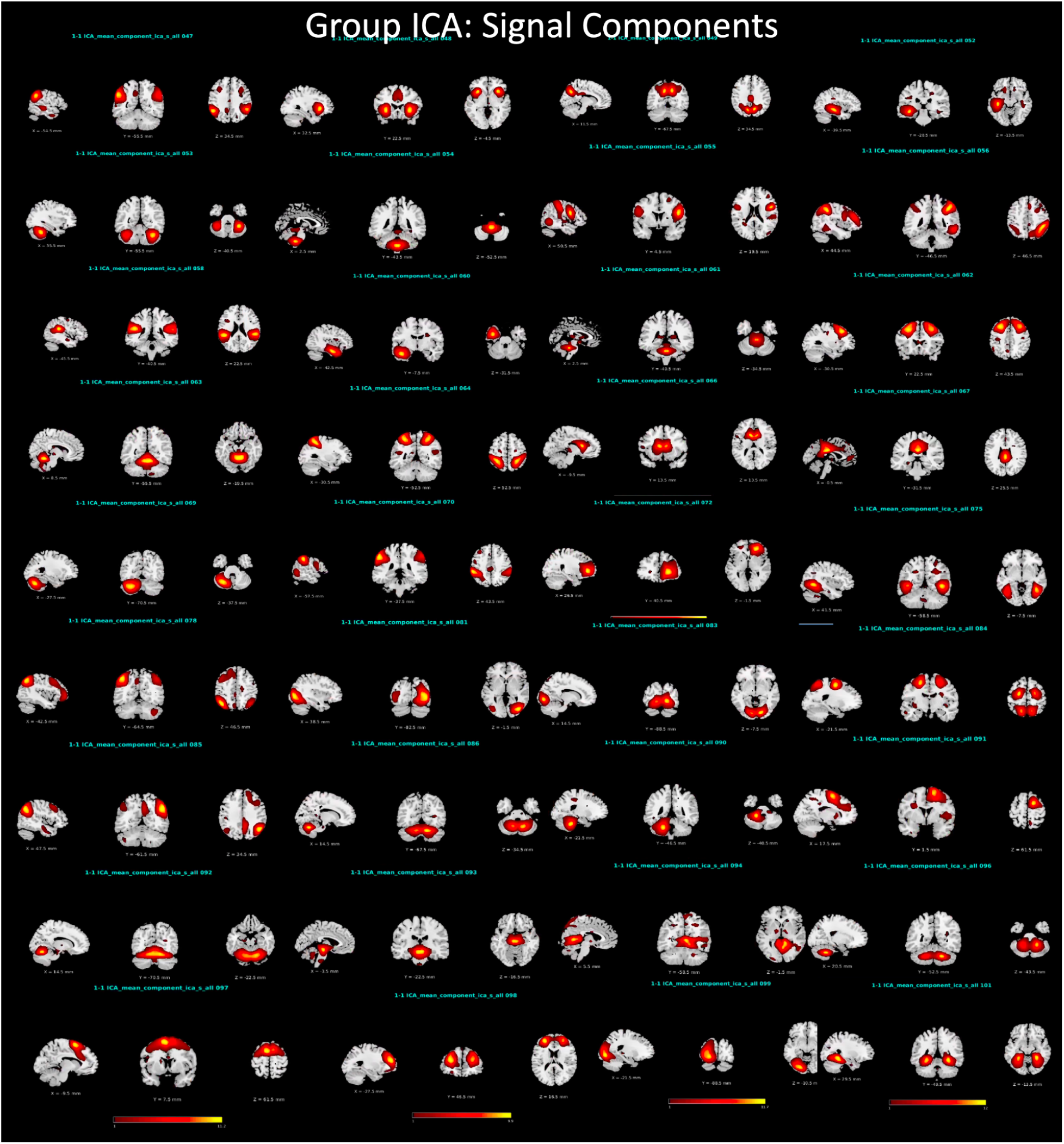
Group spatial ICA components classified as signal.

**Supplemental Figure 4:**
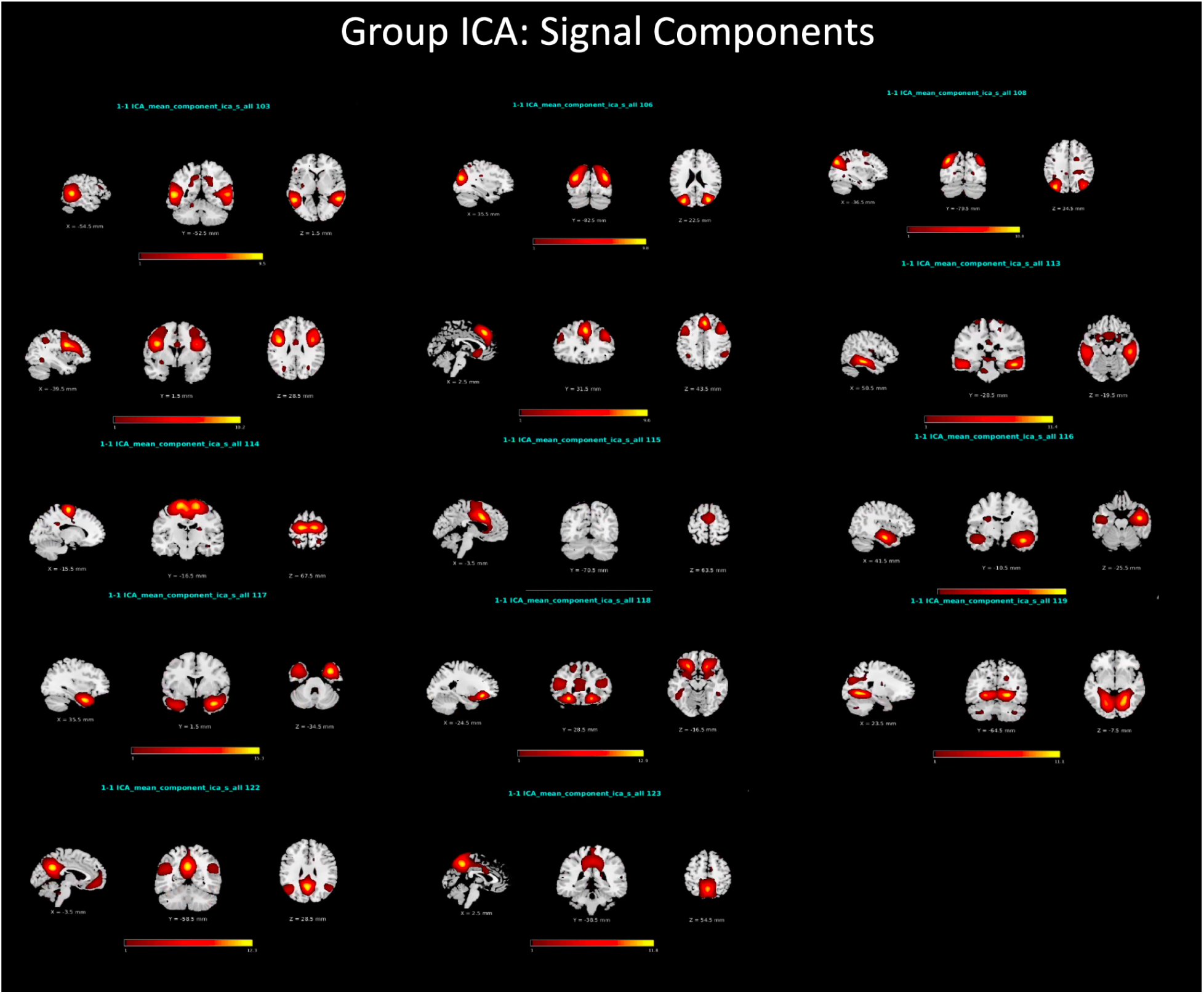
Group spatial ICA components classified as signal.

**Supplemental Figure 5:**
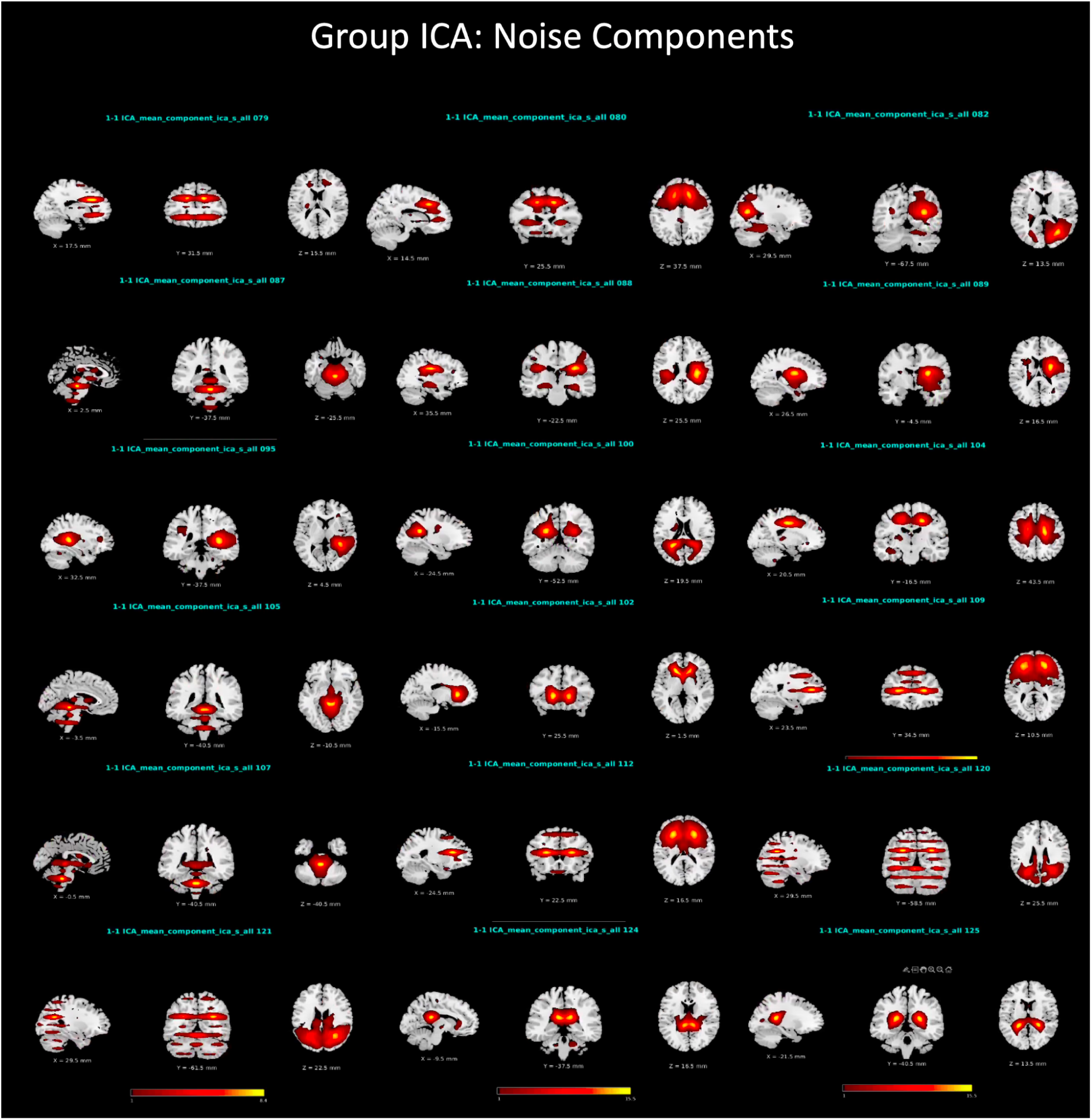
Group spatial ICA components classified as noise.

**Supplemental Figure 6:**
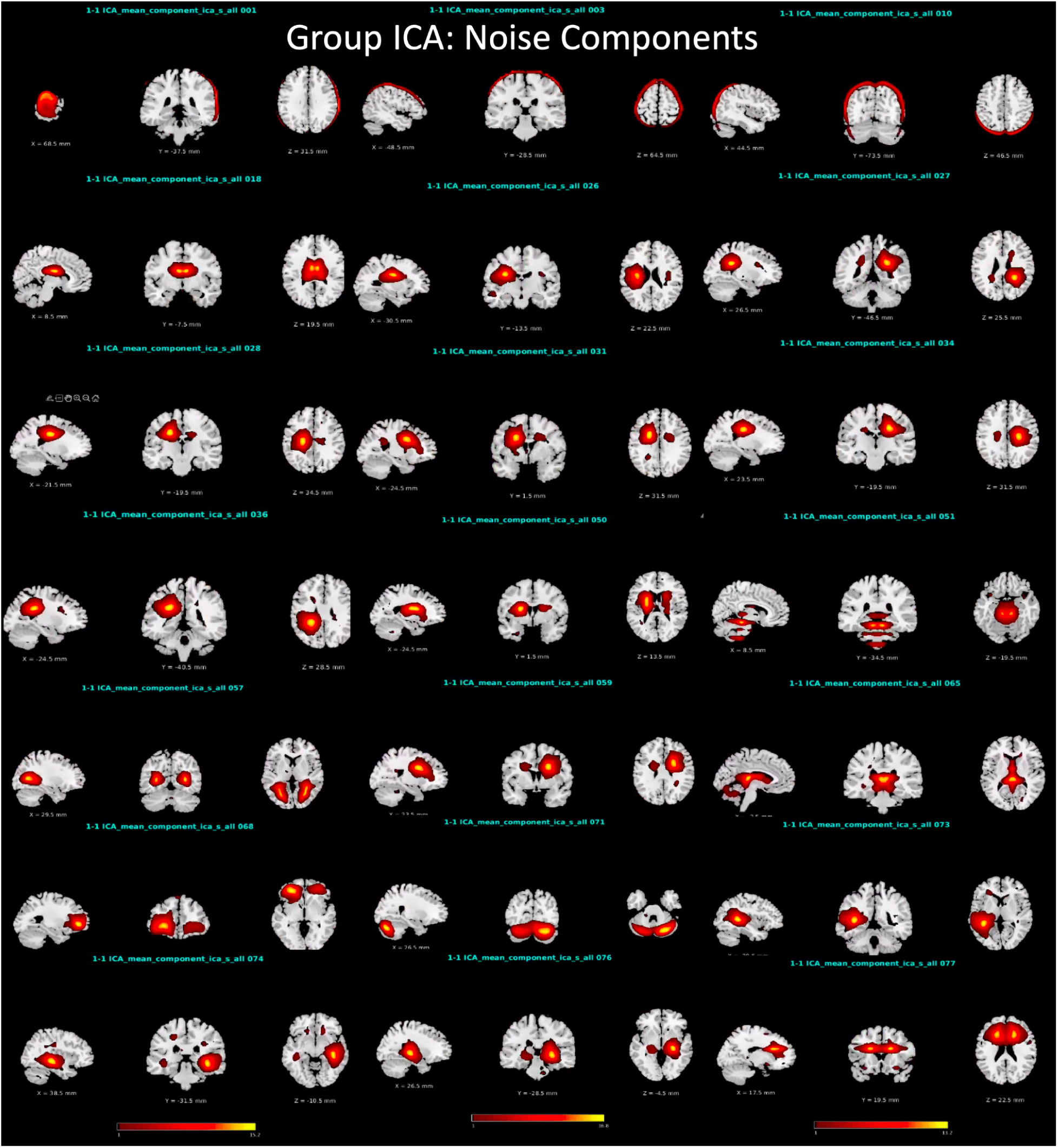
Group spatial ICA components classified as noise.

**Supplemental Figure 7:**
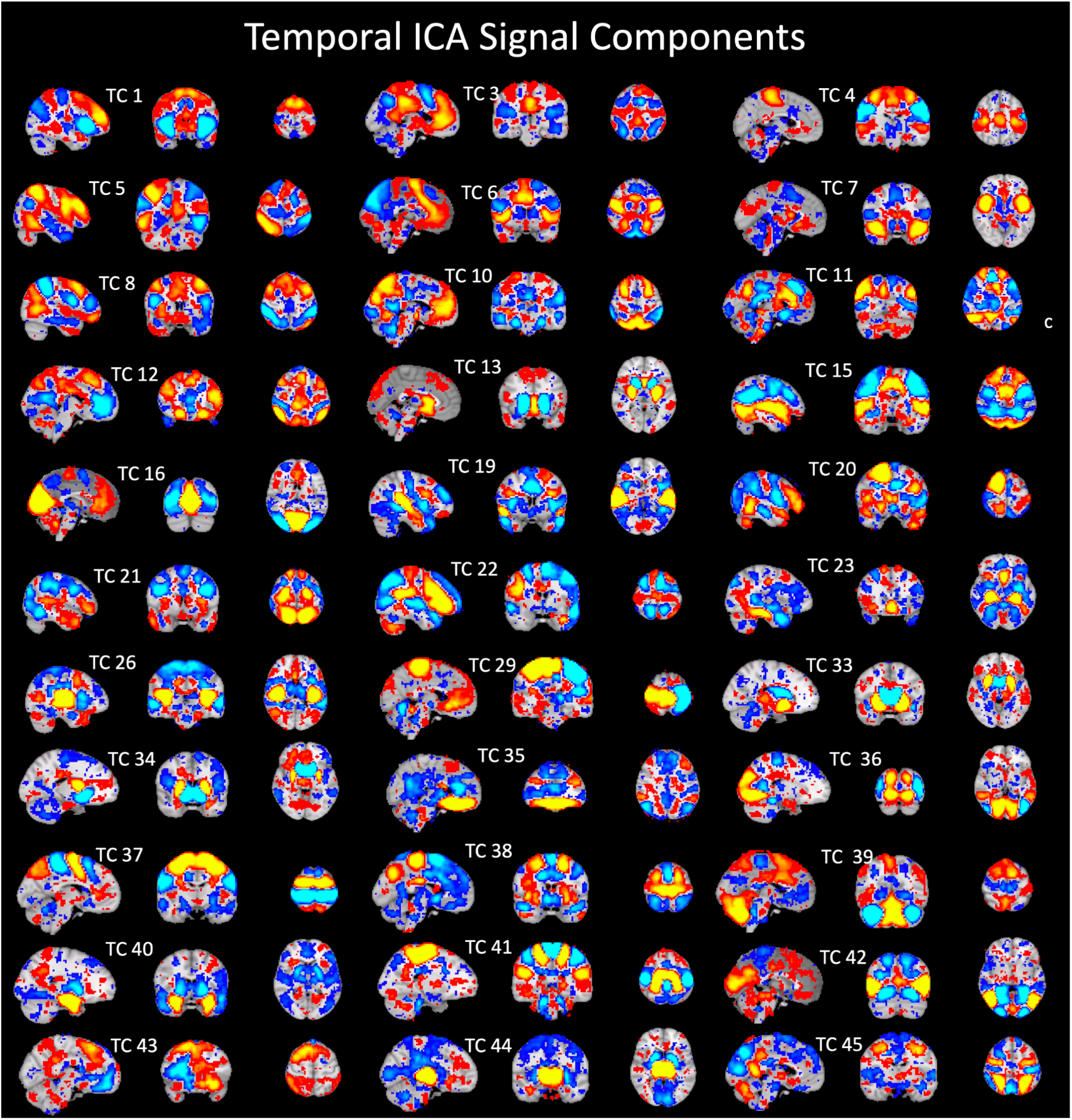
tICA components classified as signal shown with positive (hot) and negative (cold) colors.

**Supplemental Figure 8:**
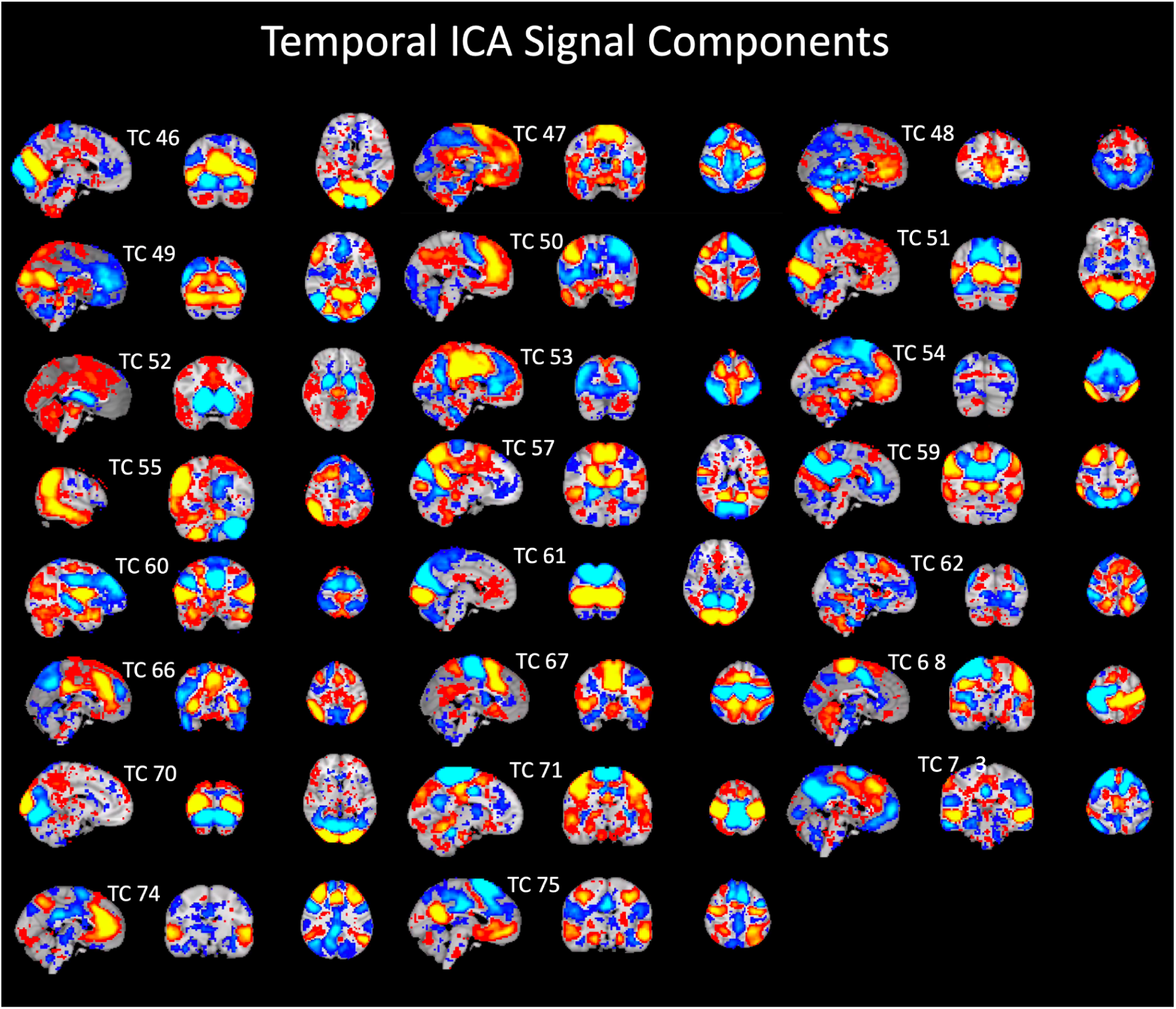
tICA components classified as neural signal shown with positive (hot) and negative (cold) colors.

**Supplemental Figure 9:**
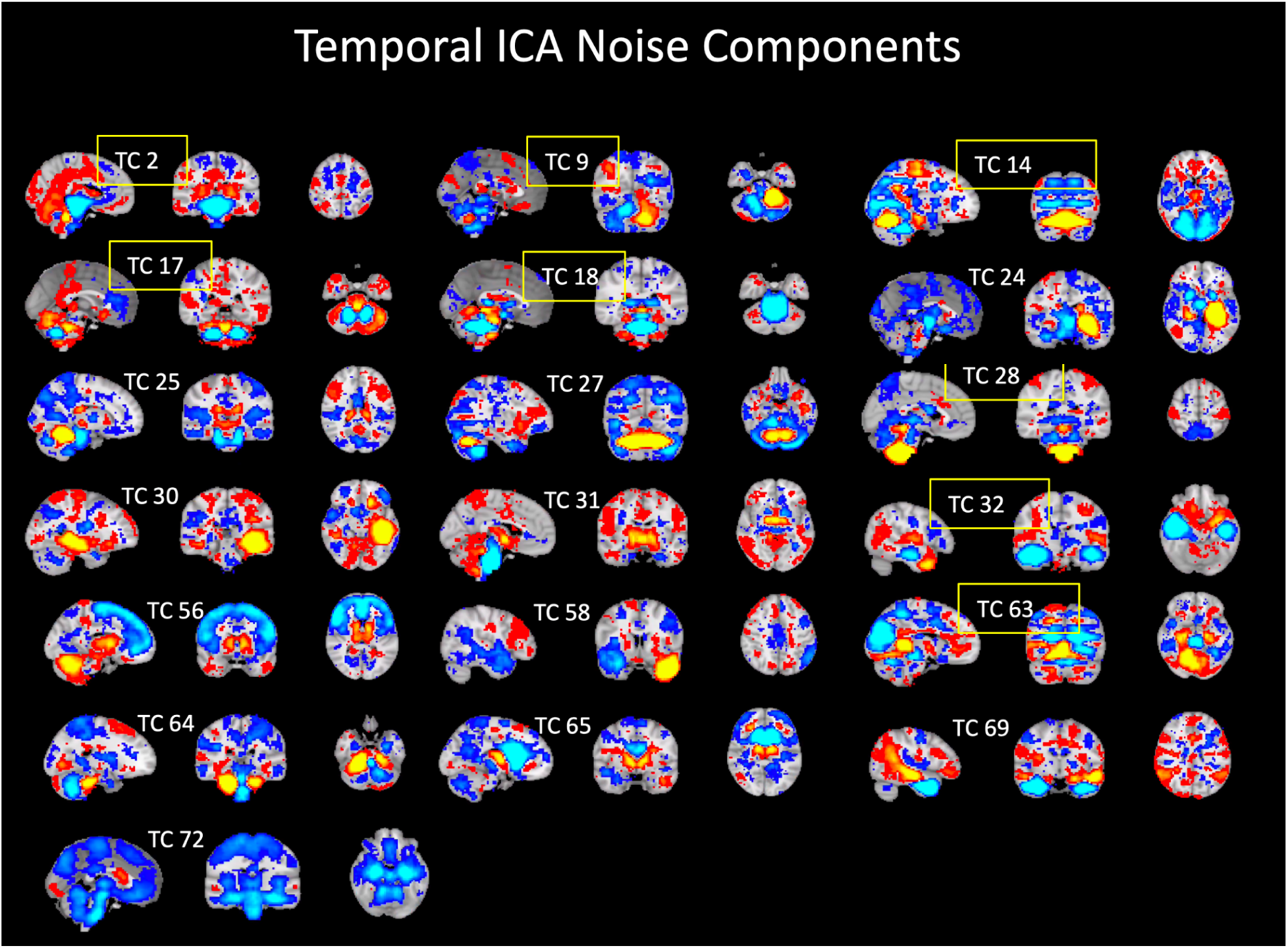
tICA components classified as noise shown with positive (hot) and negative (cold) colors. The boxes highlighted in yellow were components used in a more conservative tICA denoising with only 8 tICA noise components. The results were similar to all other pipelines. TC 72 presented a general global reduction across the brain similar to previous studies identifying this component as “global noise” (Glasser et al., 2018; Smith et al., 2012). Pipelines including or not including this component produced similar results.

**Supplemental Figure 10:**
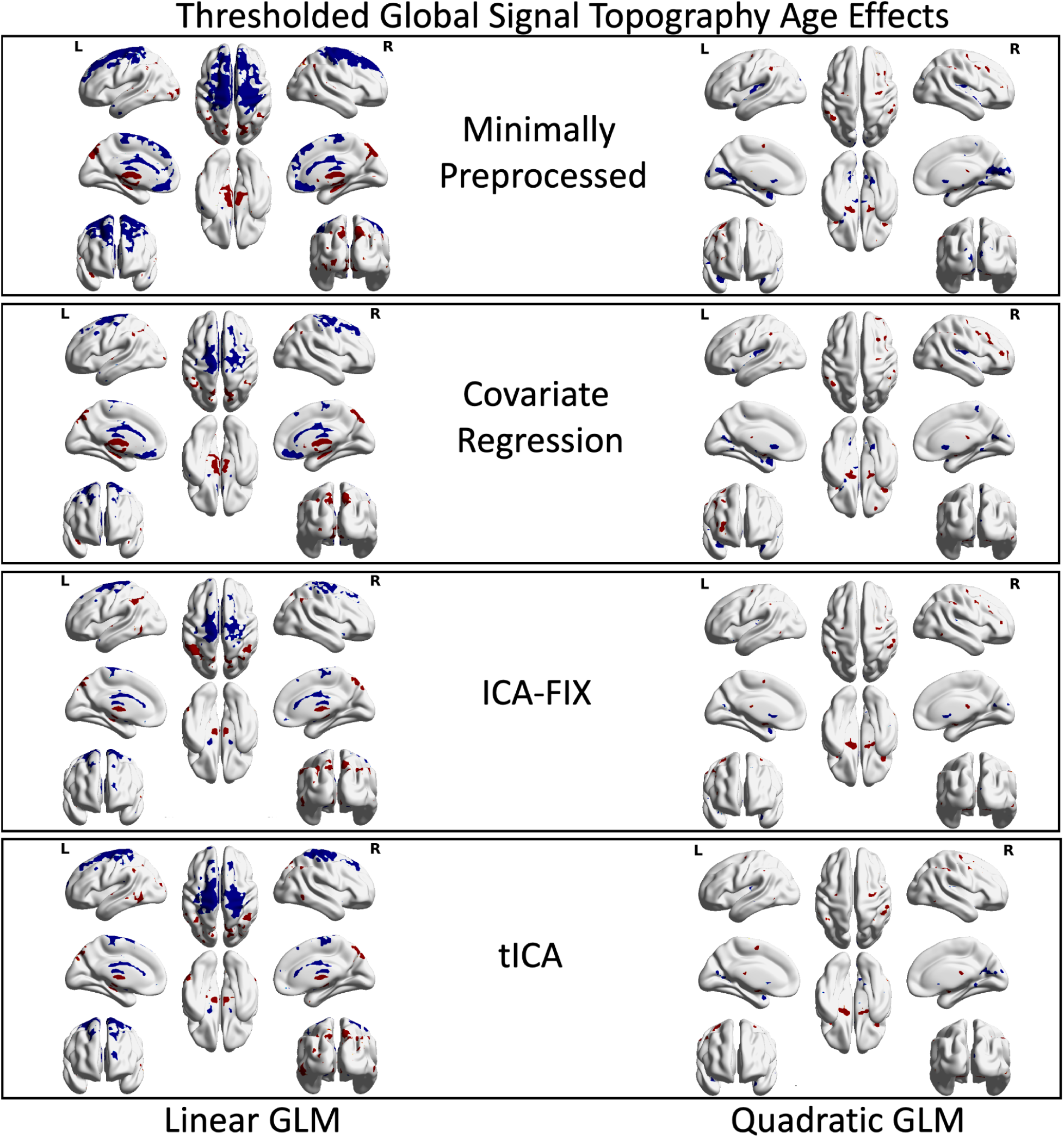
Linear and quadratic global signal topography age effect group spatial maps (voxel-wise uncorrected (*p* < 0.001) and cluster-wise corrected (*p* < 0.05). These spatial maps show the consistency of the global signal age topography effects across preprocessing pipelines. The tICA figures are the same images from main Figure 3 and are presented here for comparison.

**Supplemental Figure 11:**
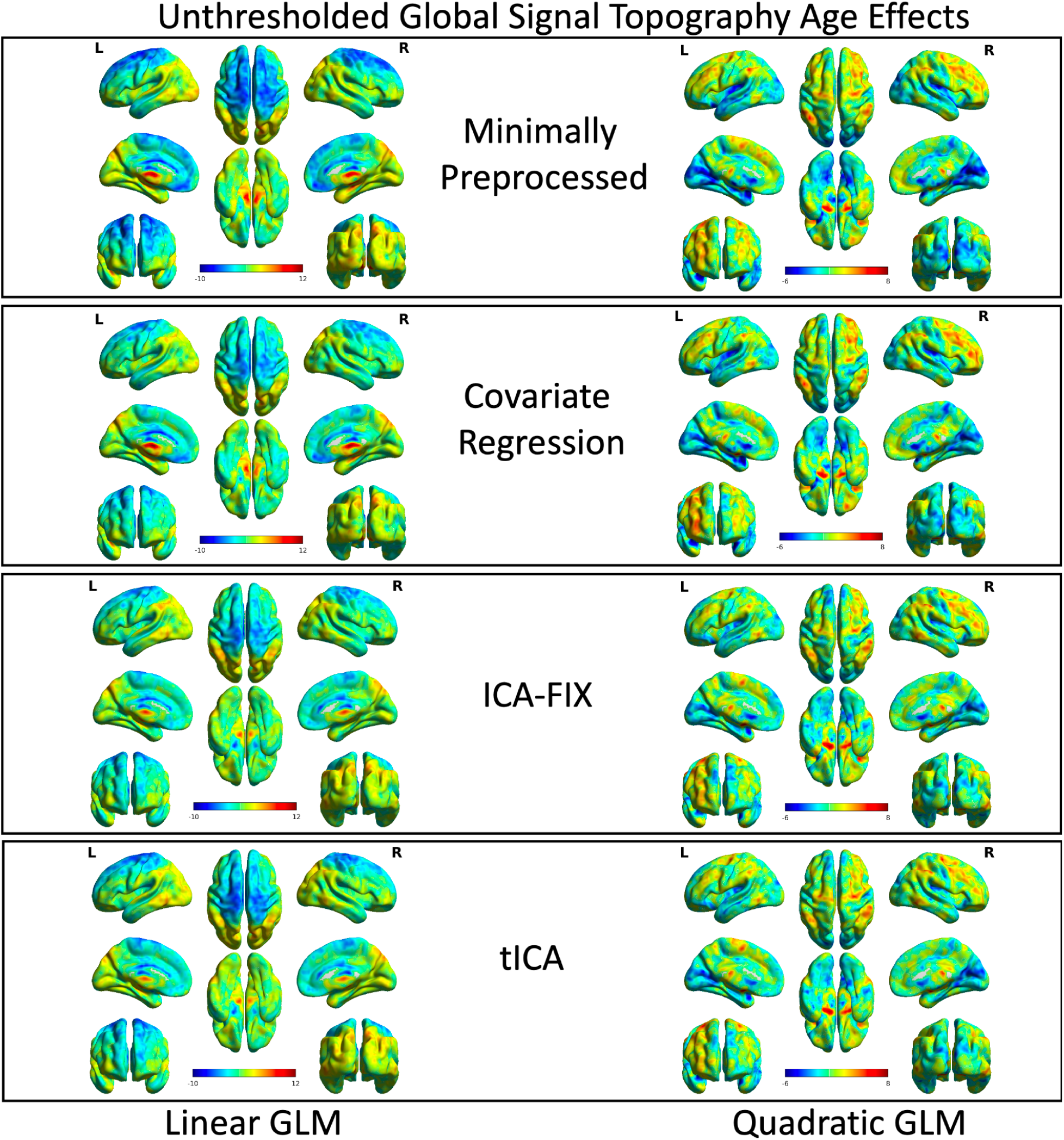
Unthresholded linear and quadratic global signal topography age effect group spatial maps. These show the consistency of the global signal age topography effects across preprocessing pipelines.

**Supplemental Figure 12:**
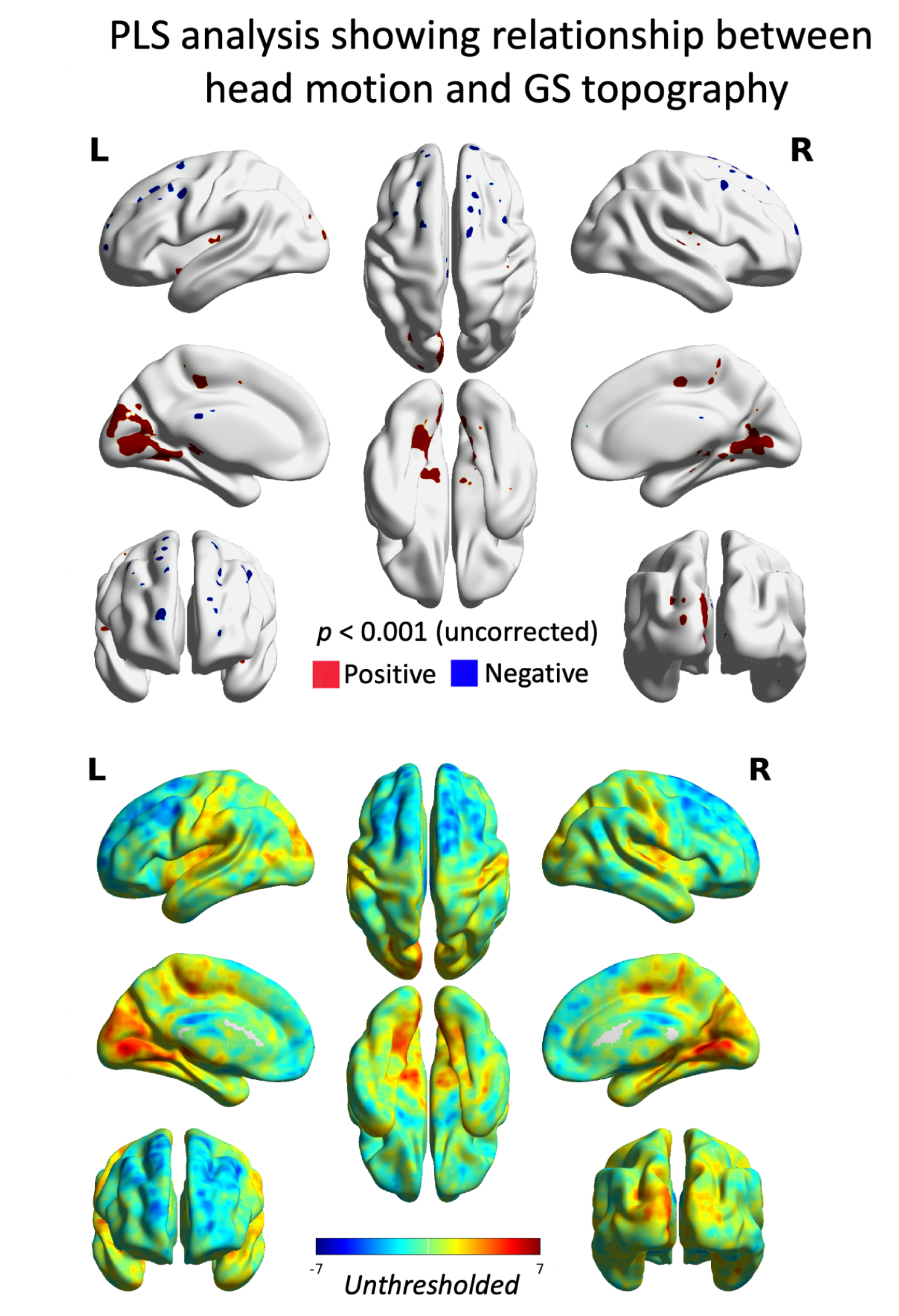
The multivariate voxel-wise spatial relationship between GS topography and head motion. A PLS analysis was conducted on individual subject GS topography spatial maps from the tICA preprocessing pipeline using framewise displacement as a behavioral head motion covariate of interest. Colors represent bootstrap ratios that approximate z values (thresholded at approximately *p* < 0.001); warmer colors represent positive associations with head motion while cooler colors represent negative associations with head motion.

**Supplemental Figure 13:**
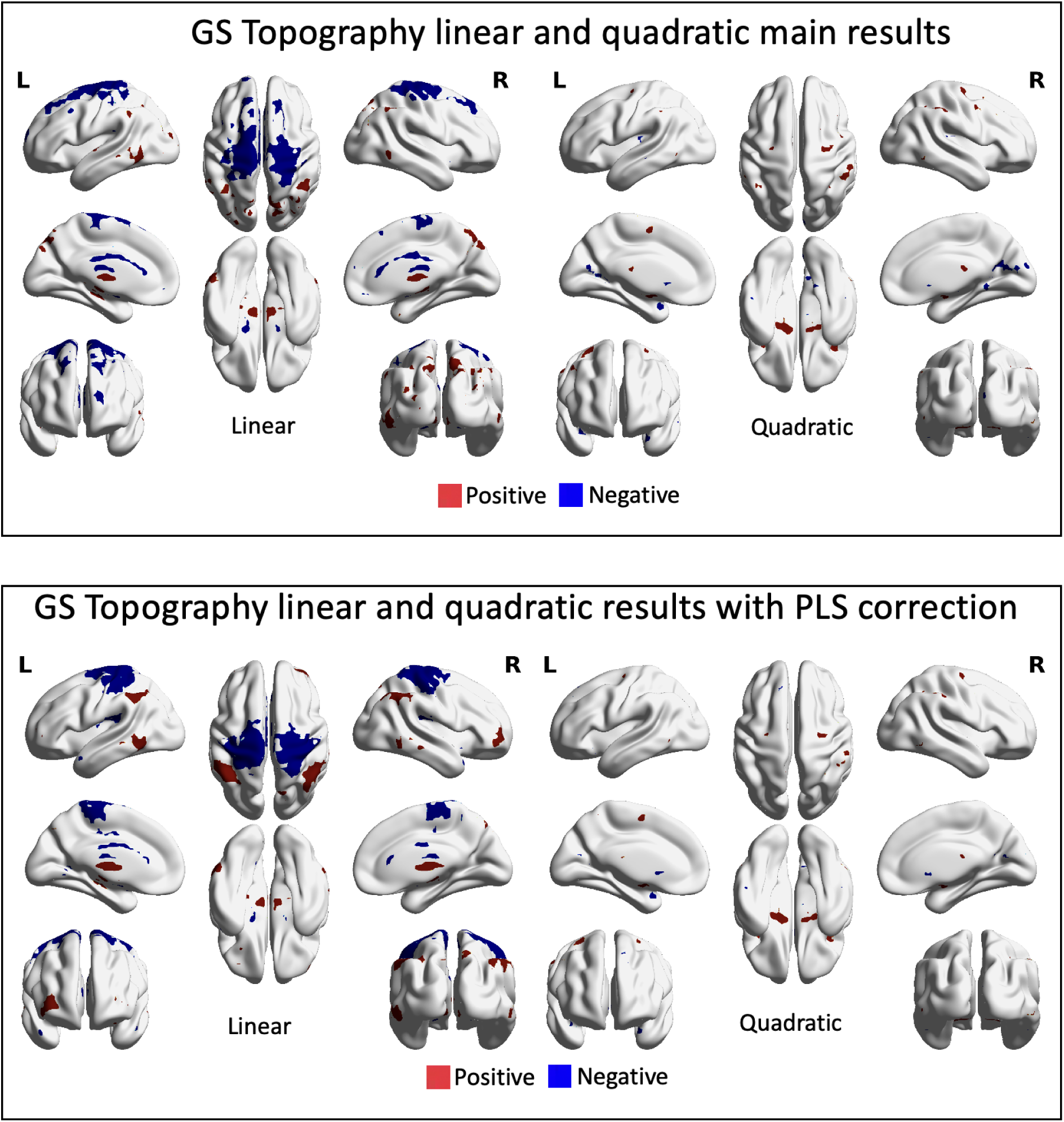
Linear and quadratic GLM results for the relationship between GS topography and age when using an additional head motion covariate from a supplemental PLS analysis. The PLS head motion covariates account for the spatial representation of head motion within individual subject GS topography maps. These analyses show that the influence of head motion is represented in a spatially distinct manner in the GS topography maps when compared to the influence of age. The top panel are the same images from main Figure 3 and are presented here for comparison purposes.

